# Oncometabolite lactate enhances breast cancer progression by orchestrating histone lactylation-dependent c-Myc expression

**DOI:** 10.1101/2023.05.14.540730

**Authors:** Madhura R. Pandkar, Sommya Sinha, Atul Samaiya, Sanjeev Shukla

## Abstract

Owing to the enhanced glycolytic rate, cancer cells generate lactate copiously, which in turn, promotes lactylation of histone. Even though histone lactylation has been explored to alter the expression of few genes, the role of this epigenetic modification in regulating the expression of oncogenes is largely unchartered. In this study, using breast cancer cell lines their mutants (which exhibit lactate-deficient metabolome), we have identified that intracellular lactate promotes histone lactylation-dependent c-Myc upregulation. Furthermore, we report that the c-Myc upregulates serine/arginine splicing factor 10 (*SRSF10*) to drive alternative splicing in breast cancer cells. Our findings provide novel mechanistic insights into the role assayed by aerobic glycolysis in orchestrating alternative splicing that collectively drive breast tumorigenesis. Moreover, we also envisage that chemotherapeutic interventions attenuating glycolytic rate can restrict breast cancer progression by impeding the c-Myc-SRSF10 axis.

## Introduction

Cancer is a malady characterized by uncontrolled proliferation of the neoplastic cells beyond the realm of normal tissue development. Therefore, cancer cells exhibit metabolic rewiring to support the requirements of enhanced replication. Seminal work performed by Otto Warburg in 1956 laid the foundation for numerous studies initiated on deciphering the altered tumor metabolism (1). It is observed that many cancer types exhibit an elevated glycolytic rate which is associated with high lactate production (2, 3). Being identified as the most commonly diagnosed type of cancer, breast cancer accounted for 11.7% of the new cancer cases in 2020, affecting over 2.3 million people world-wide (4). It is demonstrated that higher grades of breast cancers exhibit enhanced lactate levels, which makes studying the implications of lactate in breast cancer physiology indispensable (5).

Interestingly, eminent work by Zhang and co-workers recently led to the striking discovery of a previously unreported histone modification, lactylation, derived from cellular lactate (6). Although only a few reports have explored the role of histone lactylation, its contributions in regulating the expression of oncogenes that drive breast carcinogenesis strongly warrants further investigations.

Aimed to identify genes regulated by histone lactylation, in this study, we have compared the transcriptome of MCF7 (a breast cancer cell line exhibiting lactate-rich metabolome) and its PKM2 knockout mutant cells (resembling a lactate-deficient metabolome) and identified that the expression of oncogenic transcription factor c-Myc is sensitive to intracellular lactate. Our experimental investigations collectively uncovered that metabolic rewiring leading to elevated lactate production causes enrichment of histone H3 lysine 18 lactylation (H3K18la) marks on *c-Myc* promoter to result in its transcriptional upregulation. Furthermore, to elucidate the functional consequences of lactate-driven c-Myc expression, we analysed c-Myc ChIP-seq data and interestingly observed *SRSF10* as a target gene. Subsequent investigations revealed that c-Myc act as a switch that intricately modulates SRSF10 expression in response to the intracellular lactate levels. Finally, we also demonstrate that the c-Myc-SRSF10 axis regulates the alternative splicing (AS) of *MDM4* and *Bcl-x*.

Conclusively, our work provides a novel link connecting aerobic glycolysis and AS in breast cancer cells. Furthermore, we envisage that employing chemotherapeutic strategies to subside glycolytic rate can attenuate c-Myc expression to impede breast cancer progression.

## Results

### 1. Oncometabolite lactate influences breast cancer transcriptome

The rate of glycolysis-the principal producer of lactate, is essentially determined by the activity of its rate-limiting enzyme pyruvate kinase (*PKM*). Known to undergo AS, the M2 isoform of *PKM* (PKM2; found to be upregulated in various cancers), favours aerobic glycolysis. Hence, enhanced expression of PKM2 is strongly associated with higher lactate production (7). Therefore, to identify genes which are regulated in a lactate-dependent manner, we employed our previously reported model system generated in the breast cancer cells lines, where we knocked out the expression of PKM2 to establish stable lactate-deficient cell lines (8, 9) (Fig. 1A).

**Figure 1.**
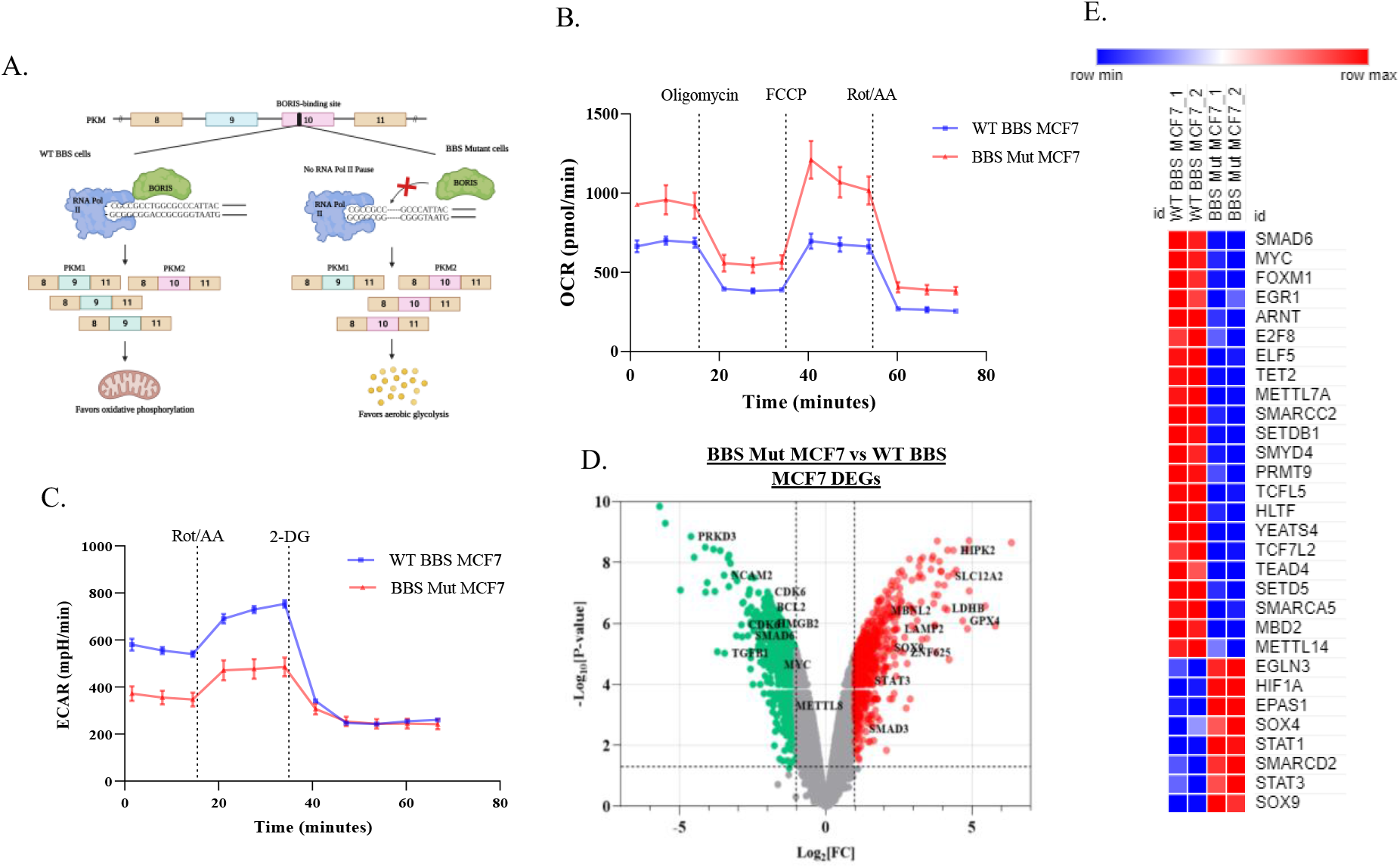
Cancer-associated metabolic affects transcriptome. *(A)* Schematic representation of CRISPR/Cas9 strategy employed to generate PKM2 knockout breast cancer cell lines. Real-time *(B)* OCR *(C) ECAR* analysis in WT BBS and BBS Mut MCF7 cells. *(D)* Volcano plot depicting DEGs in BBS Mut vs WT BBS MCF7. *(E)* Heat map representation of differentially expressed epigenetic factors and transcription factors in the BBS Mut MCF7 cells. Data are represented as mean± SD. *P ≤ 0.05.

To verify if PKM2 knockout (BBS Mut) cell lines of MCF7 and HCC1806 exhibited elevated oxidative phosphorylation (OXPHOS), we measured the oxygen consumption rate (OCR) using seahorse XFp analyzer. Our results demonstrated that the BBS Mut cells clearly exhibited higher OCR (Fig. 1B, S1.A). Moreover, the significant difference in the basal OCR (ΔOCR) denoted that due to a complete lack of PKM2 expression, BBS Mut cell lines displayed enhanced reliance on OXPHOS (Fig. S1.B). Notably, upon performing lactate assay, we observed that the WT BBS cell lines had substantially high intracellular lactate, which resulted an enhanced extracellular acidification rate (ECAR) compared to their respective BBS Mut cell lines (Fig. S1.C, 1C, S1.D). Furthermore, the difference in the basal ECAR (ΔECAR) confirmed that the WT BBS cells exhibit lactate-rich metabolome (Fig. S1.E). Collectively, our data suggests that knocking out PKM2 expression significantly decreases lactate production thereby making this model system ideal for studying the global transcriptomic alterations induced by oncometabolite lactate.

Interestingly, the analysis of Human Transcriptome Array 2.0 (HTA 2.0) data (GSE190401) of the WT BBS and BBS Mut MCF7 cells revealed that total 1962 genes (coding and non-coding) were differentially expressed in BBS Mut MCF7 cells from which, 1146 and 816 genes were downregulated and upregulated, respectively (Fig. S1.F). The significant events (P < 0.05) corresponding to the coding genes with fold change > 2 (highlighted with red points) and < -2 (highlighted with green points) are marked in figure 1D. Moreover, GO terms for biological processes enrichment analysis of differentially expressed genes (DEGs) revealed that the transcription of genes related to processes such as metabolic and cancer-related pathways, cell cycle, and spliceosome were extensively affected; underscoring the vital contributions of metabolic lactate in regulating the expression of plethora of genes involved in in diverse cancer-associated pathways (Fig. S1.G). Interestingly, numerous epigenetic factors and transcription factors identified to serve as oncogenes were differentially expressed in BBS Mut MCF7 cells (Fig. S1.H).

Conclusively, our microarray data suggests that lactate levels affect the expression of plethora of genes in breast cancer cells. Moreover, our data demonstrates that by regulating the expression of epigenetic and transcription factors, oncometabolite lactate can exert wide-reaching effects that promote breast tumorigenesis.

### 2. Elevated c-Myc expression is maintained by promoter-level histone lactylation

Strikingly, the oncogenic transcription factor c-Myc was one of the most downregulated genes in BBS Mut MCF7 cells (Fig. 1E, Table S5). Found to overexpressed in 46% of the primary breast tumors, and 30-50% of high-grade tumors, c-Myc acts as a signaling hub in regulating multiple cellular processes that drive of breast cancer progression (10–16). To investigating this intriguing possibility if c-Myc expression is responsive to metabolic lactate, we performed qRT-PCR and immunoblotting analysis and observed that c-Myc was strikingly downregulated in the BBS Mut cell lines (Fig. S2.A, 2A, S2.B). Next, as a rescue experiment, we subjected the BBS Mut cells to 15mM L-lactate treatment for 24hrs and interestingly observed heightened induction of c-Myc which confirmed that intracellular lactate drives c-Myc expression (Fig. 2B, S2.C).

**Figure 2.**
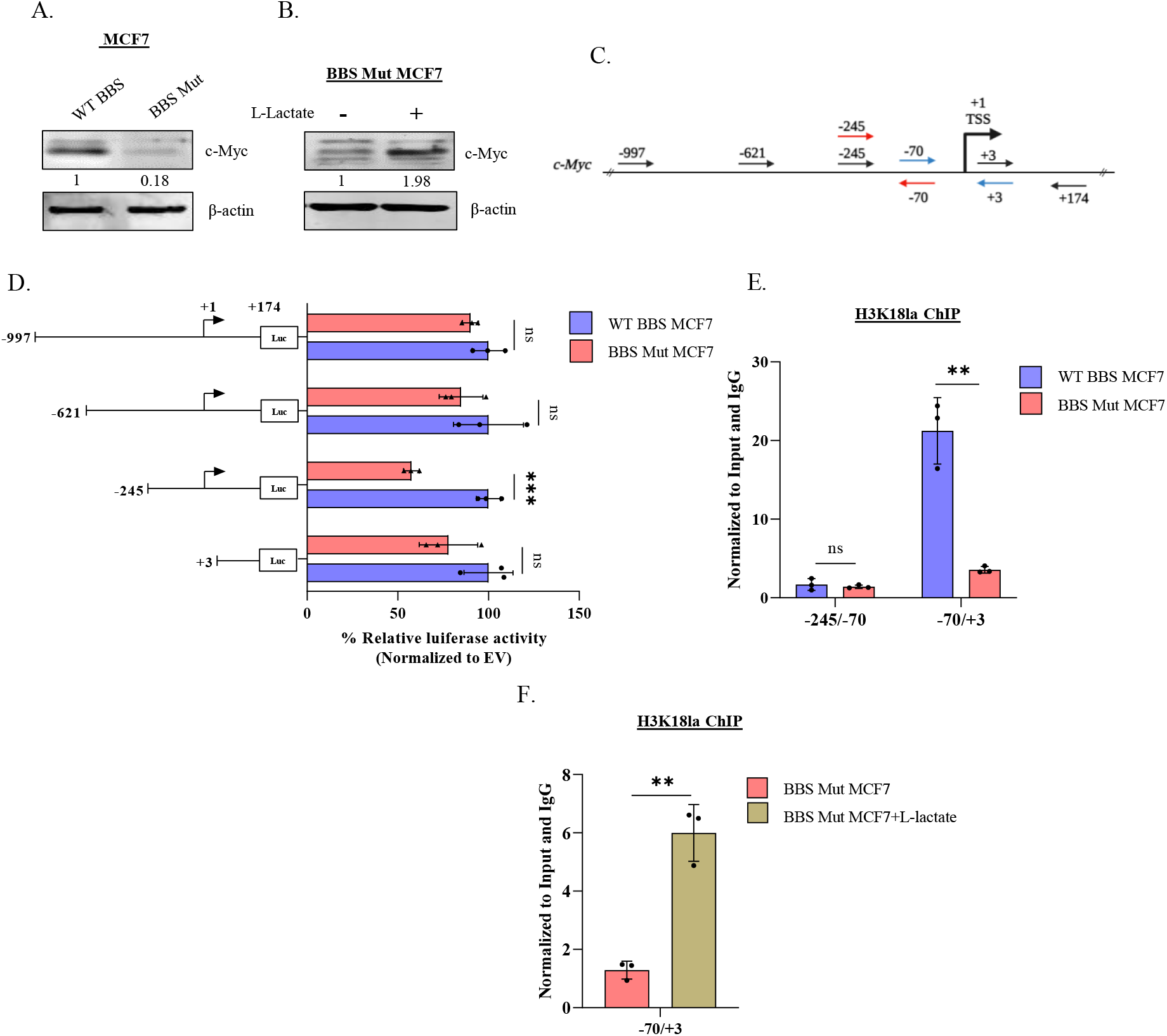
c-Myc undergoes promoter-level histone lactylation. Immunoblot analysis of c-Myc in *(A)* WT BBS and BBS Mut MCF7 cells and *(B)* on subjecting BBS Mut MCF7 cells to 15mM L-lactate treatment for 24hrs. *(C)* Schematic representation of *c-Myc* promoter. The primers used to generate deletional constructs for luciferase assay are represented using black arrows. The primers used for ChIP qRT-PCR have been marked using red and green arrows. *(D)* Luciferase assay performed using the deletional constructs of c-Myc promoter depicting a dampened activity of -245 to +174 fragment in the BBS Mut MCF7 cells. H3K18la ChIP assay performed in *(E)* WT BBS and BBS Mut MCF7 cells and *(F)* upon subjecting the BBS Mut MCF7 cells to 15mM L-lactate treatment for 12hrs. Data are represented as mean± SD. *P ≤ 0.05.

Recently, it was demonstrated that metabolic lactate can influence the histone lactylation status to regulate gene expression (6). Therefore, we analysed the data available for H3K18la ChIP-seq (GSE115354) and observed extensive enrichment of the epigenetic modification on *c-Myc* promoter (Fig. S2.D). Next, we generated multiple deletional constructs of *c-Myc* promoter for identifying the active segment primarily responsible for maintaining high c-Myc expression. Therefore, using luciferase assay, we estimated the activity of the approximately 1Kb sequence upstream of the transcriptional start site (TSS) and 174bp sequence downstream of TSS of *c-Myc* promoter (Fig. 2C). As depicted in figure 2C, a series of four deletional fragments were cloned in pGL3 basic vector: -997 to +174, -621 to +174, -245 to -174 and +3 to -174. Luciferase assay outcome demonstrated that the region between -245 to +3 exhibited decreased activity in the BBS Mut cell lines to suggest the probable involvement this region as a causative factor leading to low c-Myc expression in the BBS Mut cells (Fig. 2D, S2.E).

As p300 was previously identified as the writer of lactylation, (17, 18) we performed immunoblotting analysis of p300. No alterations in the p300 expression was detected across all the WT BBS and BBS Mut cell lines (Fig. S2.F). We also performed H3K18la chromatin immunoprecipitation (ChIP) assay to check the promoter-level histone lactylation status of the -245 to +3 region using the primers sets indicated with red and green arrows in figure 2C. Notably, the promoter region between -70 to +3 exhibited drastically decreased H3K18la marks in the BBS Mut MCF7 cells (Fig. 2E). Furthermore, upon subjecting the BBS Mut MCF7 cells L-lactate treatment, we observed a significant enrichment in H3K18la status of -70 to +3 fragment without any alterations in p300 expression (Fig. 2F, S2.G).

Collectively, our results demonstrate that the state of high intracellular lactate, as observed during elevated glycolysis, leads to H3K18la enrichment in -70 to +3 promoter region to upregulate c-Myc expression in breast cancer cells.

### 3. c-Myc regulates SRSF10 to influence alternative splicing

Next, we sought to elucidate the significance of high c-Myc expression in cancers exhibiting elevated aerobic glycolysis. Mounting evidence suggests that c-Myc has a critical role in regulating the expression of splicing factors (19–22). Moreover, our microarray results indicated that the transcription of spliceosome-related genes was affected in the lactate-deficient BBS Mut MCF7 cells (Fig. S1.G). Therefore, we hypothesized that in response to lactate, c-Myc might transcriptionally induce expression of splicing factors to function as a link between metabolism and AS. To investigate this, we analyzed c-Myc ChIP-seq data (GSM2501566) and observed its enrichment on promoter of several SRSF members; with most extensive enrichment observed on *SRSF10* promoter (Fig. 3A). Subsequently, we investigated SRSF10, as immunoblotting screen of SRSFs confirmed that only SRSF10 was downregulated in BBS Mut cells compared to its expression in the corresponding WT BBS cells, thus resembling the expression pattern of c-Myc (Fig. S3.A, S3.B).

**Figure 3.**
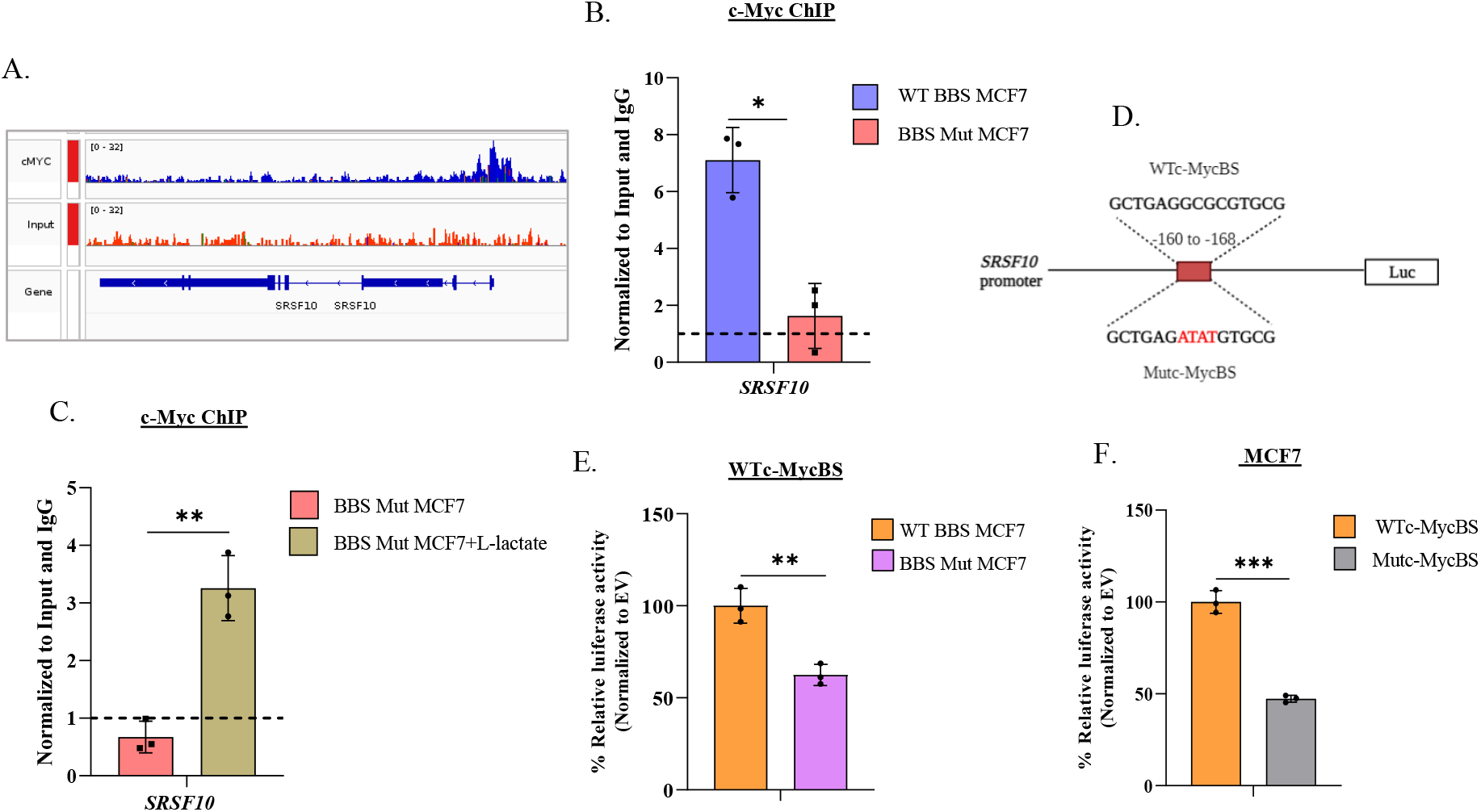
c-Myc regulates SRSF10 expression. *(A)* c-Myc ChIP-seq analysis demonstrating its enrichment on *SRSF10* promoter. c-Myc ChIP assay performed in *(B)* WT BBS and BBS Mut MCF7 cells and *(C)* upon subjecting the BBS Mut MCF7 cells to 15mM L-lactate treatment for 12hrs. *(D)* Schematic representation of *SRSF10* promoter luciferase construct depicting the wild-type c-Myc binding site (WTc-MycBS) and mutated c-Myc binding site (Mutc-MycBS). Luciferase assay of *(E)* WTc-MycBS SRSF10 construct performed in WT BBS and BBS Mut MCF7 cells and *(F)* WTc-MycBS and Mutc-MycBS SRSF10 constructs performed in WT BBS MCF7 cells. Data are represented as mean± SD. *P ≤ 0.05.

As rescue experiments, we overexpressed two critical glycolytic enzymes to enhance lactate production in BBS Mut cells. Re-introduction of PKM2 and overexpression of 6-Phosphofructo-2-Kinase/Fructose-2,6-Biphosphatase 3 (PFKFB3)-a regulator of phosphofructokinase kinase-1 (PFK-1; which catalyzes the committed step of glycolysis) stimulated c-Myc and SRSF10 expression in BBS Mut cell lines (Fig. S3.C, S3.D). Next, c-Myc knockdown caused severe reduction of SRSF10 in WT BBS cell lines (Fig. S4.E). Moreover, enforced c-Myc expression increased the endogenous SRSF10 expression in BBS Mut cell lines (Fig. S4.F). Additionally, p300 knockdown also hampered c-Myc and SRSF10 expression to confirm the H3K18la-driven regulation of c-Myc-SRSF10 axis (Fig. S3.G).

Subsequently, to identify c-Myc binding sites, we analysed *SRSF10* promoter using JASPAR database, which predicted the existence of two highly conserved overlapping c-Myc binding sites located between -160 to -168bp (23). c-Myc ChIP assay revealed its remarkable enrichment at *SRSF10* promoter in the WT BBS MCF7 cells compared to the BBS Mut MCF7 cells (Fig. 3B). Moreover, as L-lactate treatment stimulates c-Myc expression (Fig. 2B, S2.C), we performed c-Myc ChIP assay upon treating the BBS Mut MCF7 cells with 15mM L-lactate. Results clearly demonstrated an augmented c-Myc occupancy at *SRSF10* promoter in the L-lactate treated cells (Fig. 3C). Further, 113bp fragment spanning the wild-type c-Myc binding site (WTc-MycBS) present in *SRSF10* promoter was cloned upstream of a firefly luciferase (FLuc) coding sequence in pGL3 basic vector (Fig. 3D). The FLuc activity of WTc-MycBS was significantly decreased in the BBS Mut cell lines (Fig. 3E, S3.H). Next, by using site-directed mutagenesis PCR, we generated a construct of harboring mutations in the c-Myc binding site (Mutc-MycBS) (Fig. 3D). When transfected to WT BBS cells, the Mutc-MycBS construct exhibited substantially diminished FLuc activity compared to the WTc-MycBS construct to indicate c-Myc regulates SRSF10 expression in breast cancer cells (Fig. 3F, S3.I).

### 4. c-Myc-SRSF10 axis drives alternative splicing of MDM4 and Bcl-x

To understand the functional contributions of the c-Myc-SRSF10 axis, we investigated the AS events regulated by SRSF10. Previous studies have identified BCL2-like 1 (*Bcl-x*) and Murine Double Minute 4 (*MDM4*) as targets of SRSF10 (24–27). Using SpliceAid database, we identified SRSF10 binding sites on exon 9 and exon 2 of MDM4 and Bcl-x pre-mRNA respectively (Fig.S4.A, S4.B) (28), and therefore, examined the mRNA abundance of MDM4-FL isoform and the two Bcl-x isoforms (Bcl-xL and Bcl-xS) (Fig. 4A). We observed high mRNA abundance of MDM4-FL and Bcl-xL in WT BBS cells (Fig. 4B, S4.C). Furthermore, BBS Mut cell lines exhibited decreased MDM4-FL, Bcl-xL and increased Bcl-xS isoform abundance (Fig. 4B, S4.C). Notably, SRSF10 knockdown in WT BBS MCF7 cells led to decreased MDM4-FL and Bcl-xL isoforms with a concomitant enhancement in Bcl-xS isoform at mRNA level (Fig. 4C). Collectively, our results demonstrate that high SRSF10 expression supported by elevated glycolytic rate causes inclusion of *MDM4* exon 9 and favors the selection of proximal 5’ splice site of *Bcl-x*.

**Figure 4.**
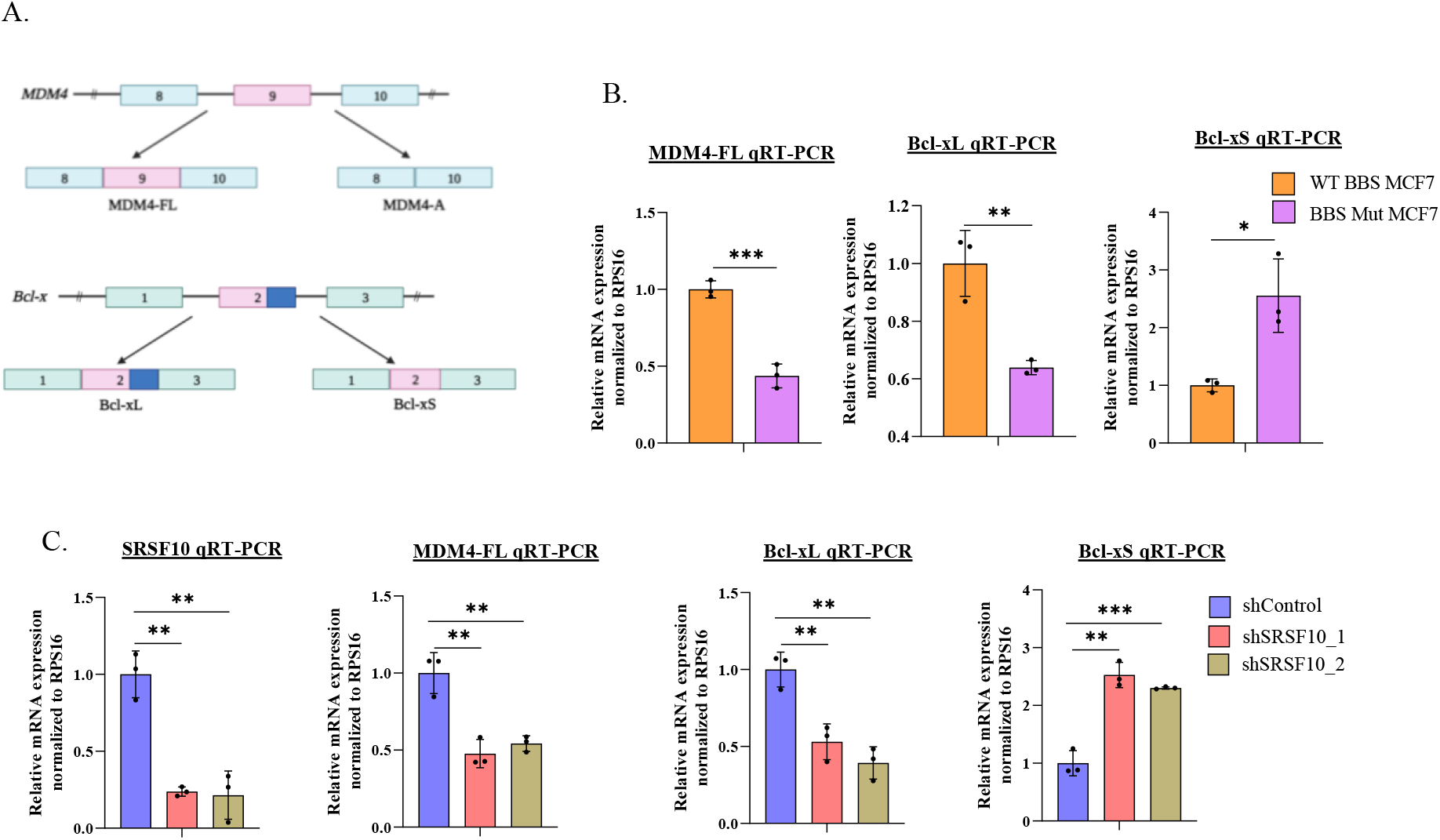
c-Myc-SRSF10 axis regulates alternative splicing of MDM4 and Bcl-x. *(A)* Schematic representation of MDM4 and Bcl-x splicing. *(B)* RPS16 normalized qRT-PCR analysis of MDM4-FL, Bcl-xL and Bcl-xS in WT BBS and BBS Mut MCF7 cells. *(C)* RPS16 normalized qRT-PCR analysis of SRSF10, MDM4-FL, Bcl-xL and Bcl-xS upon performing SRSF10 knockdown in WT BBS MCF7 cells. Data are represented as mean± SD. *P ≤ 0.05.

### 5. Targeting aerobic glycolysis restricts c-Myc-SRSF10 axis to impede proliferation of breast cancer cells

To investigate the clinical relevance of study, we examined the protein expression pattern of c-Myc and SRSF10 in breast cancer using the Clinical Proteomic Tumor Analysis Consortium (CPTAC) data. The data analysis clearly indicated that expression of c-Myc and SRSF10 is significantly higher in primary breast tumors as compared to the paired normal tissues (Fig. S5.A). Correspondingly, immunoblot analyses of 15 breast tumors samples also demonstrated a consistent upregulation of c-Myc and SRSF10 (Fig. S5.B, 5A). Furthermore, Kaplan-Meier survival analysis with The Cancer Genome Atlas (TCGA) dataset (GSE9195) showed that high c-Myc expression is associated with unfavourable patient outcome (Fig. S5.C); thus, underscoring the vitality of targeting c-Myc. Upon treating the WT BBS cells with inobrodib-a p300 inhibitor currently under clinical trial for treatment of various cancer types (29–32), we observed a significant reduction in c-Myc and SRSF10 expression (Fig. 5B). A plethora of research has highlighted targeting cancer metabolism as a desirable approach (33). Therefore, we treated the cell lines under study with a panel of metabolic inhibitors (Fig. 5C). PFK-158 and shikonin which are potent inhibitors of PFKFB3 and PKM2, respectively, are known to attenuate glycolytic rate to restrict lactate production (34, 35). PFK-158 and shikonin treatment were individually observed to downregulate c-Myc and SRSF10 expression (Fig. 5D, 5E). Demonstrated to enhance the glycolytic rate and lactate pool, rotenone treatment enhanced c-Myc and SRSF10 expression in BBS Mut cell lines (Fig. 5F) (36) .

**Figure 5.**
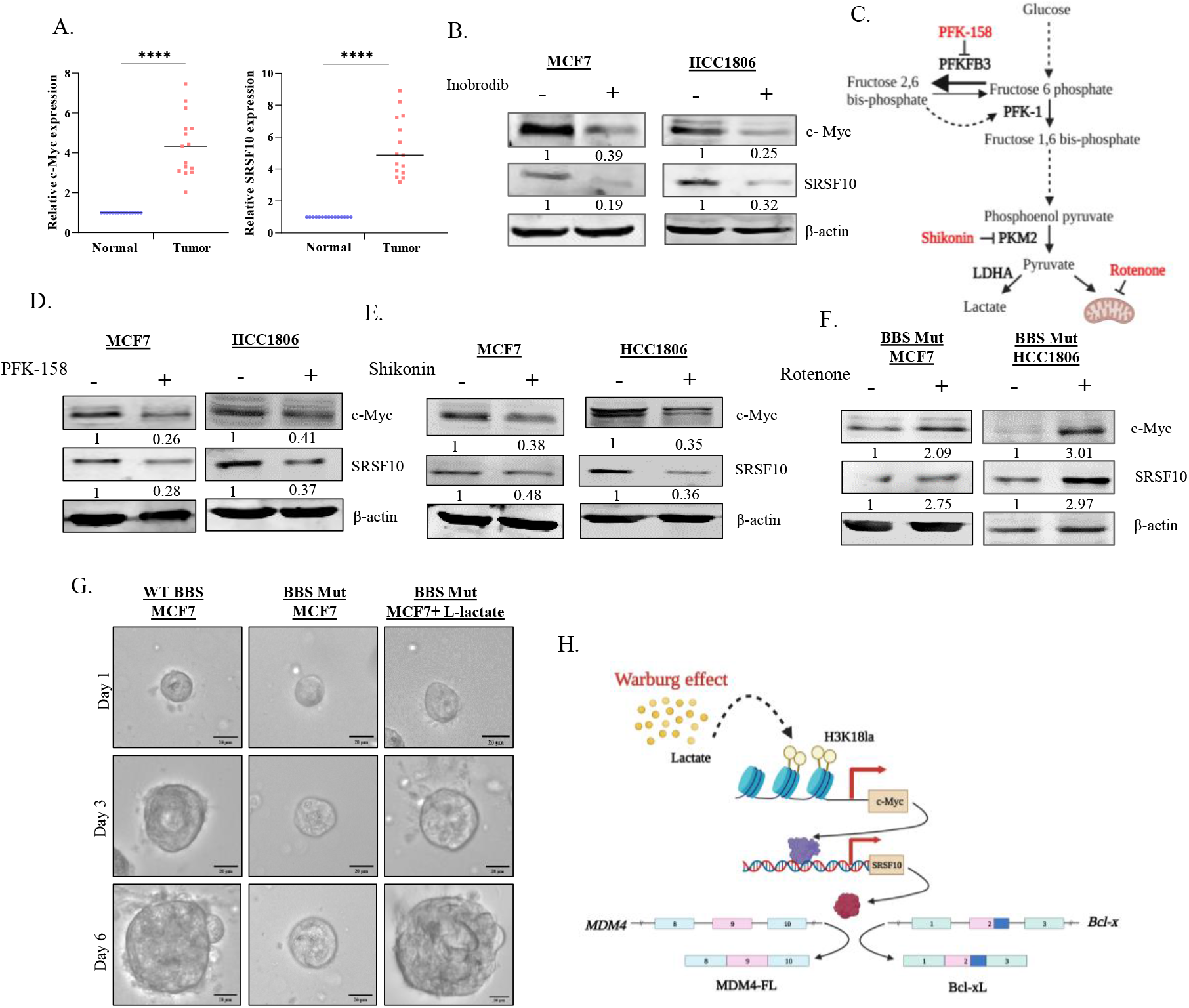
Clinical relevance of the c-Myc-SRSF10 axis. *(A)* Quantification of c-Myc and SRSF10 protein levels in 15 breast cancer patient-derived normal and tumor samples. *(B)* Immunoblot analysis showing the effect of inobrodib treatment on c-Myc and SRSF10 expression in WT BBS MCF7 cells. *(C)* Schematic representation of various metabolic inhibitors and their targets used in the study. Immunoblot analysis depicting the effect of *(D)* PFK-158 and *(E)* Shikonin treatment on c-Myc and SRSF10 expression in WT BBS MCF7 cells. *(F)* Immunoblot analysis depicting the effect of rotenone on c-Myc and SRSF10 expression in BBS Mut MCF7 cells. *(G)* Spheroid formation assay performed in WT BBS, BBS Mut, and BBS Mut MCF7 cells subjected to 5mM L-lactate treatment for 6 days *(H)* Graphical illustration of the H3K18la-drive c-Myc-SRSF10 axis. Data are represented as mean± SD. *P ≤ 0.05.

Furthermore, to evaluate if lactate-mediated upregulation of c-Myc-SRSF10 axis promoted breast carcinogenesis, we performed clonogenic assay in WT BBS, BBS Mut and BBS Mut cells cultured in media supplemented with L-lactate. Restricting lactate production by knocking out PKM2 led to substantial reduction in cell proliferation, while this phenotype was reverted upon providing L-lactate externally (Fig. S6.A, S6.B). Additionally, wound healing assay demonstrated that BBS Mut cells exhibited remarkably reduced proliferative and migratory capacity, which was rescued upon subjecting to L-lactate treatment (Fig. S6.C, S6.D). To mimic the physiological conditions in an *in-vitro* model, we monitored the growth of the spheroids derived from the WT BBS, BBS Mut and BBS Mut cell treated with L-lactate. As evident from the results, the BBS Mut cells formed remarkably smaller spheroids than their WT BBS counterpart cells. Moreover, supplementing L-lactate enhanced the proliferation of the BBS Mut cells drastically (Fig. 5G, S6.E).

Altogether, our data demonstrates that therapeutical interventions attenuating aerobic glycolysis can restrict the adverse repercussions of c-Myc-SRSF10 axis.

## Discussion

As metabolism is central to all the life processes, any metabolic abnormality results into distortion of the fine balance maintained between various cellular pathways (37). As per the findings of Otto Warburg, the arterial glucose uptake was about 47-70% and 2-18% in cancer cells and normal cells respectively; moreover, the cancer cells converted 66% of the uptaken glucose to lactate (38). Considering the vast difference in the lactate producing capabilities, it is imperative to understand the role of lactate beyond its metabolic function. The recently proposed ‘lactagenesis hypothesis’ suggests that lactate not only functions as fuel that supports expansion of cancer biomass, but also critically exhibits signaling properties, thus presenting itself as explanation and purpose of the Warburg effect in carcinogenesis (39). Consequently, there has been a growing interest in abrogating the trafficking and exchange of lactate between the cancer cells and the other neoplastic and non-neoplastic cells within the TME to restrict cancer progression (40–45).

As activation of c-Myc has been widely reported, the c-Myc driven pathways are substantially elevated in aggressive and higher grades of breast cancer. Therefore, it is imperative to identify the poorly investigated epigenetic mechanisms enabling the maintenance of high c-Myc expression. Here, in this study, we have systematically dissected the mechanism underlying oncometabolite lactate-driven epigenetic regulation of c-Myc, which provides critical evidences supporting the lactagenesis theory. Additionally, our study also highlights the role of c-Myc in regulating the expression of SRSF members. Strikingly, only SRSF10 exhibited expression pattern similar to c-Myc in response to modulations of intracellular contents to indicate that SRSF10 perhaps functions as a critical link connecting metabolism and AS (Fig.5H). Furthermore, we envisage that the MDM4 and Bcl-x splicing switch promoted by SRSF10 could have critical implications in avoidance of cellular apoptosis.

In summary, our findings uniquely demonstrate the metabolic link bridging epigenetics and oncogene-induced AS to emphasize the complexity of tumor metabolism, which warrants further studies. Moreover, our data evidently underscores the wide-reaching effects of altered metabolism that holds a paramount importance in pathogenesis of cancer.

### Experimental procedures

For detailed experimental procedures, refer to supporting information.

## Supporting information

This article contains supporting information

## Acknowledgements

The graphical representations were generated using BioRender. The authors thank all the members of Epigenetics and RNA Processing Lab, IISER Bhopal, for their critical inputs.

## Author Contributions

S. Shukla and M.R.P, conceptualization and methodology; M.R.P and S. Sinha, investigation and data analyses; S. Sinha, bioinformatic analyses; M.R.P, writing-review and editing; S. Shukla funding acquisition and supervision.

## Funding and additional information

This work is supported by Department of Biotechnology (DBT)/ Wellcome Trust India Alliance Fellowship Grant IA/I/16/2/502719, Science and Engineering Research Board (SERB) Grant (CRG/2021/004949, STR/2020/000093, IPA/2021/000148) to S. Shukla. M.R.P and S. Sinha were supported by University Grants Commission fellowship.

## References

1. O. Warburg, On the origin of cancer cells. Science (80-.). 123, 309–314 (1956).

2. R. A. Gatenby, R. J. Gillies, Why do cancers have high aerobic glycolysis? Nat. Rev. Cancer (2004) https://doi.org/10.1038/nrc1478.

3. K. G. de la Cruz-López, L. J. Castro-Muñoz, D. O. Reyes-Hernández, A. García-Carrancá, J. Manzo-Merino, Lactate in the Regulation of Tumor Microenvironment and Therapeutic Approaches. Front. Oncol. (2019) https://doi.org/10.3389/fonc.2019.01143.

4. H. Sung, et al., Global Cancer Statistics 2020: GLOBOCAN Estimates of Incidence and Mortality Worldwide for 36 Cancers in 185 Countries. CA. Cancer J. Clin. 71, 209–249 (2021).

5. S. M. Cheung, et al., Lactate concentration in breast cancer using advanced magnetic resonance spectroscopy. Br. J. Cancer 2020 1232 123, 261–267 (2020).

6. D. Zhang, et al., Metabolic regulation of gene expression by histone lactylation. Nature 574, 575–580 (2019).

7. F. Guillaumond, et al., Strengthened glycolysis under hypoxia supports tumor symbiosis and hexosamine biosynthesis in pancreatic adenocarcinoma. Proc. Natl. Acad. Sci. U. S. A. 110, 3919–3924 (2013).

8. M. R. Pandkar, et al., PKM2-mediated epigenetic reprogramming regulates hypoxic expression of PFKFB3 to promote breast cancer progression. bioRxiv, 2022.11.06.515384 (2022).

9. S. Singh, et al., Intragenic DNA methylation and BORIS-mediated cancer-specific splicing contribute to the Warburg effect. Proc. Natl. Acad. Sci. U. S. A. 114, 11440–11445 (2017).

10. P. Chrzan, J. Skokowski, A. Karmolinski, T. Pawelczyk, Amplification of c-myc gene and overexpression of c-Myc protein in breast cancer and adjacent non-neoplastic tissue. Clin. Biochem. 34, 557–562 (2001).

11. X. Zhang, et al., Mechanistic insight into Myc stabilization in breast cancer involving aberrant Axin1 expression. Proc. Natl. Acad. Sci. U. S. A. 109, 2790–2795 (2012).

12. E. Yeh, et al., A signalling pathway controlling c-Myc degradation that impacts oncogenic transformation of human cells. Nat. Cell Biol. 2004 64 6, 308–318 (2004).

13. J. Blancato, B. Singh, A. Liu, D. J. Liao, R. B. Dickson, Correlation of amplification and overexpression of the c-myc oncogene in high-grade breast cancer: FISH, in situ hybridisation and immunohistochemical analyses. Br. J. Cancer 2004 908 90, 1612–1619 (2004).

14. S. L. Deming, S. J. Nass, R. B. Dickson, B. J. Trock, C-myc amplification in breast cancer: a meta-analysis of its occurrence and prognostic relevance. Br. J. Cancer 2000 8312 83, 1688–1695 (2000).

15. M. Eilers, R. N. Eisenman, Myc’s broad reach. Genes Dev. 22, 2755–2766 (2008).

16. C. V. Dang, MYC on the Path to Cancer. Cell 149, 22–35 (2012).

17. Z. Yang, et al., Lactylome analysis suggests lactylation-dependent mechanisms of metabolic adaptation in hepatocellular carcinoma. Nat. Metab. 2023 51 5, 61–79 (2023).

18. J. Xiong, et al., Lactylation-driven METTL3-mediated RNA m6A modification promotes immunosuppression of tumor-infiltrating myeloid cells. Mol. Cell 82, 1660-1677.e10 (2022).

19. W. Zhao, et al., Splicing factor derived circular RNA circUHRF1 accelerates oral squamous cell carcinoma tumorigenesis via feedback loop. Cell Death Differ. 2019 273 27, 919–933 (2019).

20. C. Caggiano, M. Pieraccioli, V. Panzeri, C. Sette, P. Bielli, c-MYC empowers transcription and productive splicing of the oncogenic splicing factor Sam68 in cancer. Nucleic Acids Res. 47, 6160–6171 (2019).

21. C. J. David, M. Chen, M. Assanah, P. Canoll, J. L. Manley, HnRNP proteins controlled by c-Myc deregulate pyruvate kinase mRNA splicing in cancer. Nature (2010) https://doi.org/10.1038/nature08697.

22. L. Urbanski, et al., MYC regulates a pan-cancer network of co-expressed oncogenic splicing factors. Cell Rep. 41, 111704 (2022).

23. O. Fornes, et al., JASPAR 2020: update of the open-access database of transcription factor binding profiles. Nucleic Acids Res. 48, D87–D92 (2020).

24. X. Zhou, et al., Transcriptome analysis of alternative splicing events regulated by SRSF10 reveals position-dependent splicing modulation. Nucleic Acids Res. 42, 4019–4030 (2014).

25. M. Sohail, et al., A novel class of inhibitors that target SRSF10 and promote p53-mediated cytotoxicity on human colorectal cancer cells. NAR cancer 3, 1–22 (2021).

26. S. Yadav, et al., ERK1/2-EGR1-SRSF10 Axis Mediated Alternative Splicing Plays a Critical Role in Head and Neck Cancer. Front. Cell Dev. Biol. 9, 2468 (2021).

27. H. Li, et al., SRSF10 Regulates Alternative Splicing and Is Required for Adipocyte Differentiation. https://doi.org/10.1128/MCB.01674-13 34, 2198–2207 (2023).

28. F. Piva, M. Giulietti, L. Nocchi, G. Principato, SpliceAid: a database of experimental RNA target motifs bound by splicing proteins in humans. Bioinformatics 25, 1211–1213 (2009).

29. L. Nicosia, et al., Potent Pre-Clinical and Early Phase Clinical Activity of EP300/CBP Bromodomain Inhibitor CCS1477 in Multiple Myeloma. Blood 140, 852–853 (2022).

30. L. Nicosia, et al., Therapeutic Targeting of EP300/CBP By Bromodomain Inhibition in Acute Myeloid Leukemia. Blood 140, 8774–8775 (2022).

31. J. Welti, et al., Targeting the p300/cbp axis in lethal prostate cancer. Cancer Discov. 11, 1118–1137 (2021).

32. L. Helminen, et al., Chromatin Accessibility and Pioneer Factor FOXA1 Shape Glucocorticoid Receptor Action in Prostate Cancer. bioRxiv, 2023.03.03.530941 (2023).

33. Z. E. Stine, Z. T. Schug, J. M. Salvino, C. V. Dang, Targeting cancer metabolism in the era of precision oncology. Nat. Rev. Drug Discov. 2021 212 21, 141–162 (2021).

34. Y. Xiao, et al., Inhibition of PFKFB3 induces cell death and synergistically enhances chemosensitivity in endometrial cancer. Oncogene 2021 408 40, 1409–1424 (2021).

35. J. Chen, et al., Shikonin and its analogs inhibit cancer cell glycolysis by targeting tumor pyruvate kinase-M2. Oncogene 30, 4297–4306 (2011).

36. W. L. Hou, et al., Inhibition of mitochondrial complex I improves glucose metabolism independently of AMPK activation. J. Cell. Mol. Med. 22, 1316 (2018).

37. M. R. Pandkar, S. G. Dhamdhere, S. Shukla, Oxygen gradient and tumor heterogeneity: The chronicle of a toxic relationship. Biochim. Biophys. Acta - Rev. Cancer 1876, 188553 (2021).

38. O. Warburg, F. Wind, E. Negelein, THE METABOLISM OF TUMORS IN THE BODY. J. Gen. Physiol. 8, 519 (1927).

39. I. San-Millán, G. A. Brooks, Reexamining cancer metabolism: lactate production for carcinogenesis could be the purpose and explanation of the Warburg Effect. Carcinogenesis 38, 119–133 (2017).

40. T. Fiaschi, et al., Reciprocal metabolic reprogramming through lactate shuttle coordinately influences tumor-stroma interplay. Cancer Res. 72, 5130–5140 (2012).

41. P. Sanità, et al., Tumor-stroma metabolic relationship based on lactate shuttle can sustain prostate cancer progression. BMC Cancer 14, 1–14 (2014).

42. P. Sonveaux, et al., Targeting lactate-fueled respiration selectively kills hypoxic tumor cells in mice. J. Clin. Invest. 118, 3930–3942 (2008).

43. F. Morais-Santos, et al., Targeting lactate transport suppresses in vivo breast tumour growth. Oncotarget 6, 19177 (2015).

44. N. Draoui, et al., Synthesis and pharmacological evaluation of carboxycoumarins as a new antitumor treatment targeting lactate transport in cancer cells. Bioorg. Med. Chem. 21, 7107–7117 (2013).

45. J. R. Doherty, J. L. Cleveland, Targeting lactate metabolism for cancer therapeutics. J. Clin. Invest. 123, 3685 (2013).

