## Supporting information for "Oncometabolite lactate enhances breast cancer progression by orchestrating histone lactylation-dependent c-Myc expression"

**Running title:** c-Myc is epigenetically upregulated by histone-lactylation

**Keywords:** Epigenetics, breast cancer, Warburg effect, histone modifications, tumor metabolism, alternative splicing, c-Myc, SRSF10.

### Supporting Information

### Experimental procedures

#### 1. Cell culture

Human breast cancer cell lines MCF7 and HCC1806 were procured from American Type Culture Collection (ATCC). MCF7 and HEK293T were cultured at 37ºC, 5% $CO_2$ in DMEM (Invitrogen, 12800017, lot no. 2248833). RPMI-1640 (Invitrogen, 23400021, lot number 2144859) was used to culture HCC1806. The culture media for all the cell lines was supplemented with 10% fetal bovine serum (FBS; Sigma, F7524, lot. no BCBX8466), 100 units/ml of penicillin and streptomycin (Invitrogen, 15140122, lot no. 2321120), and 2mM/L L-glutamine (Invitrogen, 25030081, lot no. 1917006). The BBS Mut MCF7 and HCC1806 cell lines were generated as previously stated (8, 9). The media and cell culture conditions for various CRISPR/Cas9 mutants were identical to the wild-type cells.

#### 2. Molecular cloning

The PKM2 and PFKFB3 overexpression constructs were generated as previously mentioned (8, 9). Briefly, full length PKM2 was amplified using MCF7 cDNA as template and cloned between NotI and SalI sites of the plasmid pCMV-3tag-1a. Similarly, full length PFKFB3 was amplified using MCF7 cDNA as template and cloned between NotI and EcoRI sites of pCMV-3tag-1a plasmid. Full length c-Myc was also amplified from MCF7 cDNA and cloned between BamHI and EcoRI sites in pCMV-3tag-1a. The sequence of the obtained construct was verified using Sanger sequencing. The details of the primers used are provided in the table S2.

#### 3. Luciferase reporter assay

*c-Myc* promoter fragments and *SRSF10* promoter fragment were individually amplified using MCF7 genomic DNA as template and subcloned between the KpnI and HindIII sites of pGL3 basic vector (Promega, E1751). The sequence of the obtained constructs was verified using Sanger sequencing. The details of the primers used for generating luciferase constructs are enlisted in the table S2. Cells were seeded in a 24-well plate and allowed to attach overnight.

The wells were co-transfected with different *c-Myc* or *SRSF10* promoter constructs along with pRL-TK renilla luciferase plasmid. 24hrs post transfection, the cells were lysed and luciferase activity was determined. The relative luciferase activity was determined by dividing the firefly luciferase activity with the renilla luciferase activity.

**4. Site-directed mutagenesis**

The site-directed mutant construct of *SRSF10* promoter was generated using oligonucleotides harboring desired mutations in the c-Myc binding site located from -160bp to -168bp relative to the TSS. The wild-type *SRSF10* promoter construct was used as template. The sequence of the SDM construct was verified by Sanger sequencing. The primers used for obtaining SDM construct are enlisted in table S2. A similar protocol as mentioned for luciferase assay was followed for transfecting and obtaining readings of the SDM construct.

**5. Immunoblotting**

The cells were lysed in urea lysis buffer (8M urea, 2M thiourea, 2% CHAPS, 1% DTT) supplemented with 1× protease inhibitor cocktail (PIC; leupeptin 10-100M, pepstatin 1M, EDTA 1-10mM, AEBSF 1mM) at 4°C for 30min and centrifuged for 2 hours at maximum speed (16,900g). The supernatant was separated, quantified, and an equal concentration of protein samples was loaded. After separation proteins were electro-transferred on an active PVDF membrane. Following transfer, the blots were incubated overnight at 4°C with suggested dilutions of primary antibodies, followed by 1 hour incubation with secondary antibody. The Odyssey membrane Scanning equipment was used to scan the blots. The bands were quantified using GelQuant software (version 1.8.2). The details of the antibodies used are provided in table S1.

**6. Quantitative RT-PCR**

Total RNA was extracted according to the manufacturer's instructions using TRIZol reagent (Ambion, 15596018, lot no. 260712). The concentration was measured using an Eppendorf BioSpectrometer, and 2µg of total RNA was reverse transcribed using the Invitrogen SuperScript® III First-Strand Synthesis System (18080-051, lot no. 2291381). Amplifications were carried out in duplicates using the GO taq QPCR master mix (Promega, A6002, lot no. 0000385100) and the Roche light cycler 480 II according to the manufacturer's instructions. Primers were designed using IDT PrimerQuest tool (https://www.idtdna.com/) and are listed in table S3. Using the formula $2^{\wedge(Ct\_control-Ct\_target)}$, the average cycle thresholds from three independent biological replicates were computed and normalised to the housekeeping control gene RPS16. The Student's t-test was performed to compare gene/exon expression levels between two groups. P< 0.05 was considered statistically significant.

### 7. Chromatin Immunoprecipitation (ChIP) assay

The ChIP assay was performed as previously mentioned (8). Briefly, 10 approximately 10 million cells were crosslinked, scaped in PBS, lysed and sonicated. 25μg of sheared chromatin was immunoprecipitated with an antibody of interest following overnight incubation at 4°C. The immunoprecipitated protein-DNA complexes and 5% input were purified to eliminate proteins and the eluted DNA was analysed by qRT-PCR using GO taq QPCR master mix (Promega, A6002, lot no. 0000385100) in triplicate using primers specific for *c-Myc* or *SRSF10* promoter. Each experiment was performed atleast thrice and normalizations were performed using the formula $2^{\wedge(Ct\ input\ -\ Ct\ immunoprecipitation)}$. The obtained values were normalized to relative rabbit IgG and control IP values. The primers were designed using IDT PrimerQuest tool (https://www.idtdna.com/) and are listed in table S3. Significance between the two groups was calculated using Student's t-test, with a value of <0.05 considered statistically significant.

### 8. RNA interference

$3\times10^5$ cells were seeded per well of a six-well culture plate and allowed to attach for 24hrs. The lentivirus containing small hairpin RNA (shRNA) (Sigma, Mission Human Genome shRNA Library) against the target gene was inoculated in the presence of 8μg/ml polybrene (Sigma, H9268, lot no. SLBH5907V) containing media. Cells were selected for 72hrs using 1μg/ml puromycin (Sigma, P9620, lot no. 034M4008V) and subsequently used for various experiments. The sequence of shRNAs used in this study is listed in table S4.

### 9. Lactate assay

Following treatment, an equal number of WT BBS and BBS Mut MCF7 and HCC1806 cells were lysed using ice cold assay buffer provided in the lactate assay kit (Sigma, MAK064-1, lot no. 3F09K06270). Lactate quantification was carried out using the deproteinized lysates according to the manufacturer's instructions. The readings were taken at room temperature with a microplate reader set to 450nm optical density. Student's t-test was used to calculate statistical significance.

### 10. Extracellular flux assays

Oxygen consumption rate (OCR), extracellular acidification rate (ECAR) was determined using seahorse XF HS mini analyzer. $5 \times 10^3$ cells per well were seeded and allowed to attach overnight in an 8-well seahorse XFp mini cell culture plate. Cells were then washed and incubated with XF assay medium supplemented with 1mM pyruvate (Sigma, S8636, lot no. RNBJ2351), 2mM L-glutamine (Invitrogen, 25030081, lot no. 1917006), 10mM glucose (Gibco, A24940-01, lot no. 1969830) and incubated 1hr at 37°C in CO2-free incubator. OCR and ECAR estimation was performed as per manufacturer’s instructions. OCR was assessed in response to oligomycin (1.5µM), FCCP (0.5µM), and rotenone/antimycin A (0.5µM) and ECAR was assessed in response to rotenone/antimycin A (0.5µM), and 2-deoxy-D-glucose (2-DG; 50mM). Finally, the readings were normalized to the respective protein concentrations.

### 11. Clonogenic assay

$5 \times 10^3$ cells were seeded in each well of a 6-well cell culture plate and were cultured for 10 days. The media was replaced every after 72hrs. The cells washed thrice with 1X PBS and were then fixed using 4% formaldehyde for 15min at RT. The cells were then stained with 0.05% crystal violet solution prepared in 10% ethanol. The cells were then gently washed thrice with 1X PBS and the plates were air dried for 15 min and imaged.

**12. Wound healing assay**

$3\times10^5$ cells were seeded in 6-well plate. Upon reaching confluency, a 200µl sterile pipette tip was used to create a wound. The plate was then washed twice with 1X PBS to remove debris. The images of same region was captured at 0, 24 and 48hrs with an inverted microscope.

**13. Generation of spheroid cultures**

The spheroids were generated as previously described (8). Briefly, 50µL of Growth Factor Reduced (GFR) Basement Membrane Matrix (Corning, 356230, lot no. 9343006) was spread in a well of 96-well cell culture plate and allowed to solidify at 37°C for 1hr. $7\times10^3$ cells were resuspended in 100µL of cell-culture media supplemented with 1µg/mL hydrocortisone (Sigma, H0888, lot no. SLBG4963V) and 5µg/mL insulin (Sigma, I1882, lot no. SLBR1114V) and added over the solidified matrix. Media containing 10% GFR matrigel was then subsequently overlayed as the topmost layer and incubated at 37°C in the presence of 5% $CO_2$. The media containing hormones was replaced every 72hrs and images were captured using Thermo Scientific EVOS FL Auto 2 imaging system.

**14. Breast cancer sample collection**

Tumor and adjacent normal tissue pairs were collected from patients undergoing surgery for breast cancer at Bansal Hospital, Bhopal, India. For the study, approval was granted by Institute Ethics Committee of Indian Institute of Science Education and Research Bhopal. Informed consent was obtained from all the patients. The tissue samples were snap frozen immediately after surgery and stored at −80°C until use. Clinical characteristics of patients used in the study are provided in Table S6.

**15. Human Transcriptome Array 2.0**

HTA 2.0 was performed as previously described (8). WT BBS and BBS Mut MCF7 cells were harvested using TRIZol reagent (Ambion, 15596018, lot no. 260712) and RNA was isolated using PureLink RNA Mini Kit (Invitrogen, 12183025, lot no. 1862249) as per manufacturer's instructions. RNA concentration was estimated using Eppendorf BioSpectrometer and as per manufacturer’s protocol, 100ng of total RNA was used for biotinylated cDNA synthesis using

GeneChip™ WT Plus Reagent Kit (Invitrogen, 902281, lot no. 01059768). Subsequently, cDNA fragmentation was performed, and 5.5µg of fragmented cDNA was hybridized on Affymetrix GeneChip™ Human Transcriptome Array 2.0 (HTA2.0) chips (Invitrogen, 902162, lot no. 4364589) for 16 hrs at 45°C. The chips were washed and stained using the Affymetrix Fluidics Station 450. Post hybridization, the fluorescence intensity of the arrays was scanned using the Affymetrix Scanner 7G. The raw files generated in CEL format were used for further analysis. The microarray data generated in this study is deposited under the GEO accession number: GSE190401.

**16. Human Transcriptome Array 2.0 data analysis**

The CEL files were analyzed using Transcriptome Array Console 4.0 (Invitrogen, version 4.0.2.15) using the gene+exon–SST-RMA method of summarization. Genes with thresholds of absolute fold-change >2 and < -2, P<0.05, and false discovery rates (FDRs) <0.05 were selected as differentially expressed genes (DEGs). The volcano plot for DEGs and violin plot for differentially expressed epigenetic factors and transcription factors were generated using GraphPad Prism 9 software. A heat map for differentially expressed epigenetic factors and transcription factors was generated using online tool Morpheus (https://software.broadinstitute.org/morpheus) and the over-representation analysis (ORA) for GO biological processes was generated using ShinyGo 0.76.2 (45). FDR <0.05 was considered significant while obtaining enriched GO terms.

**17. CPTAC data analysis**

Protein expression profile of c-Myc and SRSF10 and MYC was analysed in CPTAC breast cancer dataset in normal vs primary tumor tissues using University of Alabama Cancer (UALCAN) platform (46).

**18. Survival data analysis**

Recurrence-free survival of c-Myc was analyzed in cohort GSE9195 using online tool Kaplan–Meier Plotter (www.kmplot.com). The best possible cut-off was used as upper and lower quartile. Samples were divided into high- and low-expression groups and compared for recurrence-free survival.

### 19. ChIP-seq data analyses

The available ChIP-seq data for H3K18la (GSE115354) and c-Myc (GSM2501566) were downloaded from Gene Expression Omnibus database. The raw fastQ files were downloaded and trimmed by Trimmomatic (v0.39) with default parameters. STAR (v2.7.3a) aligner was used to uniquely align the reads to GRCh38.Peak calling was done using MACS2 v2.1.2.

### 20. Statistical analysis

GraphPad Prism 9 was used for all statistical analyses. Unless otherwise specified, all data are presented as mean ±SD and were analysed using a two-tailed Student's t-test. The statistical methods for each analysis are provided in the figure legends or in the materials and methods sections. P-values less than 0.05 were deemed significant. ns = not significant, *P ≤ 0.05, **P ≤ 0.01, ***P ≤ 0.001, ****P ≤ 0.0001.

### Supporting information (S1)

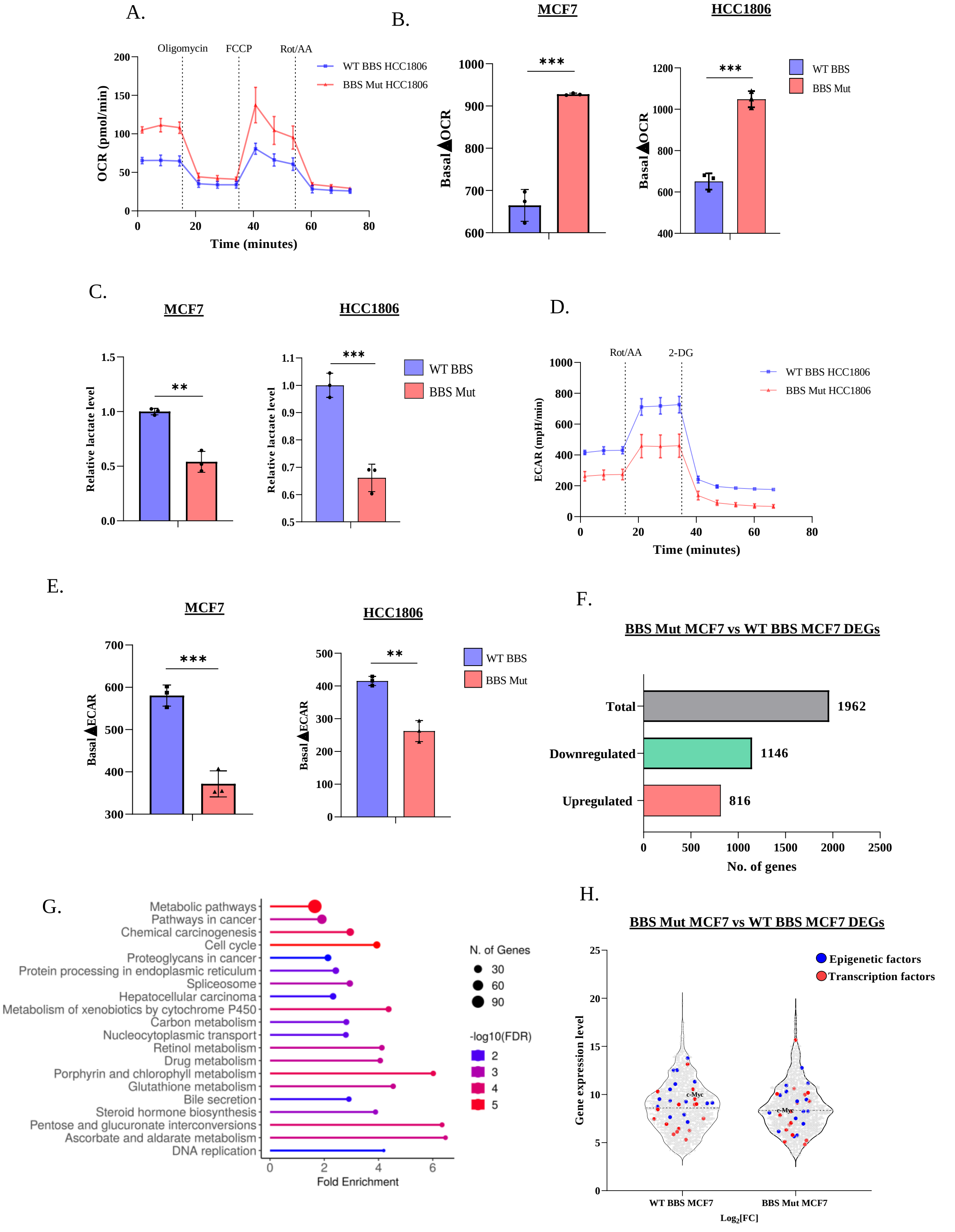


**Aerobic glycolysis affects transcriptome.** *(A)* Real-time OCR analysis performed in WT BBS and BBS Mut HCC1806 cells. *(B)* Quantification of differential basal OCR (ΔOCR) in the WT BBS and BBS Mut cell lines. *(C)* Intracellular lactate production in WT BBS and BBS Mut cell lines estimated by lactate assay. *(D)* Real-time ECAR analysis in WT BBS and BBS Mut HCC1806 cells. *(E)* Quantification of differential basal ECAR (ΔECAR) in the WT BBS and BBS Mut cells. *(F)* Bar graph representing DEGs in BBS Mut vs WT BBS MCF7 cells using HTA 2.0 data. *(G)* The top 20 enriched terms for biological processes for the DEGs obtained from the microarray data. *(H)* Violin plot depicting the differentially expressed epigenetic factors (blue points) and transcription factors (red points) in BBS Mut MCF7 cells. Data are represented as mean± SD. *$P \leq 0.05$.

### Supporting information 2 (S2)

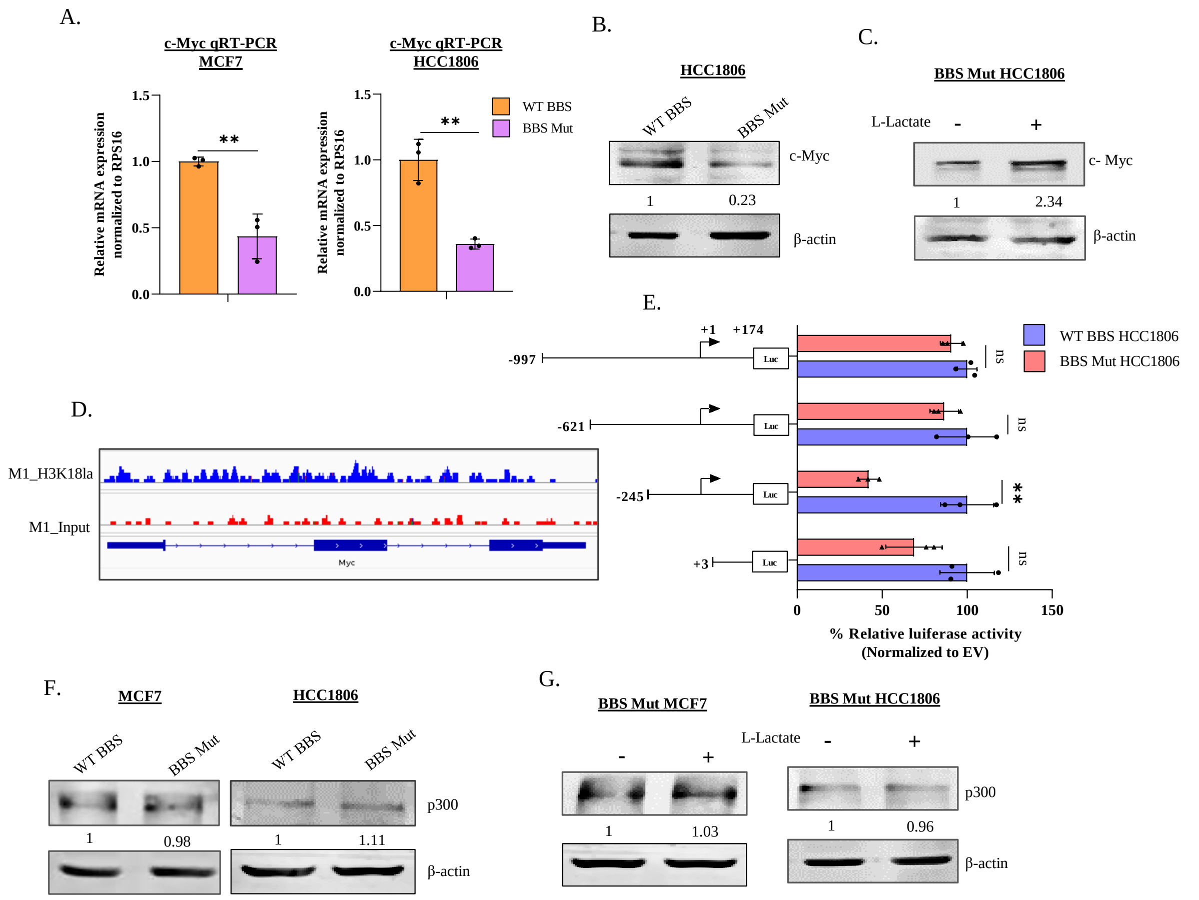


**Intracellular lactate regulates transcription of c-Myc**. *(A)* RPS16 normalized qRT-PCR analysis of *c-Myc* in WT BBS and BBS Mut cell lines. *(B)* Immunoblot analysis depicting a reduced expression of c-Myc in BBS Mut HCC1806 cells. *(C)* Immunoblot analysis of c-Myc on subjecting BBS Mut HCC1806 cells to 15mM L-lactate treatment for 24hrs. *(D)* H3K18la ChIP-seq analysis indicating enrichment of the histone marks on c-Myc promoter. *(E)* Luciferase assay performed using the deletional constructs of *c-Myc* promoter depicting a dampened activity of -245 to +174 fragment in the BBS Mut HCC1806 cells. *(F)* Immunoblot analysis of p300 in WT BBS and BBS Mut cell lines. Results demonstrate insignificant change in p300 expression. *(G)* Immunoblot analysis of p300 upon subjecting the BBS Mut cells to 15mM L-lactate treatment for 24hrs. Data are represented as mean± SD. *P ≤ 0.05.

### Supporting information 3 (S3)

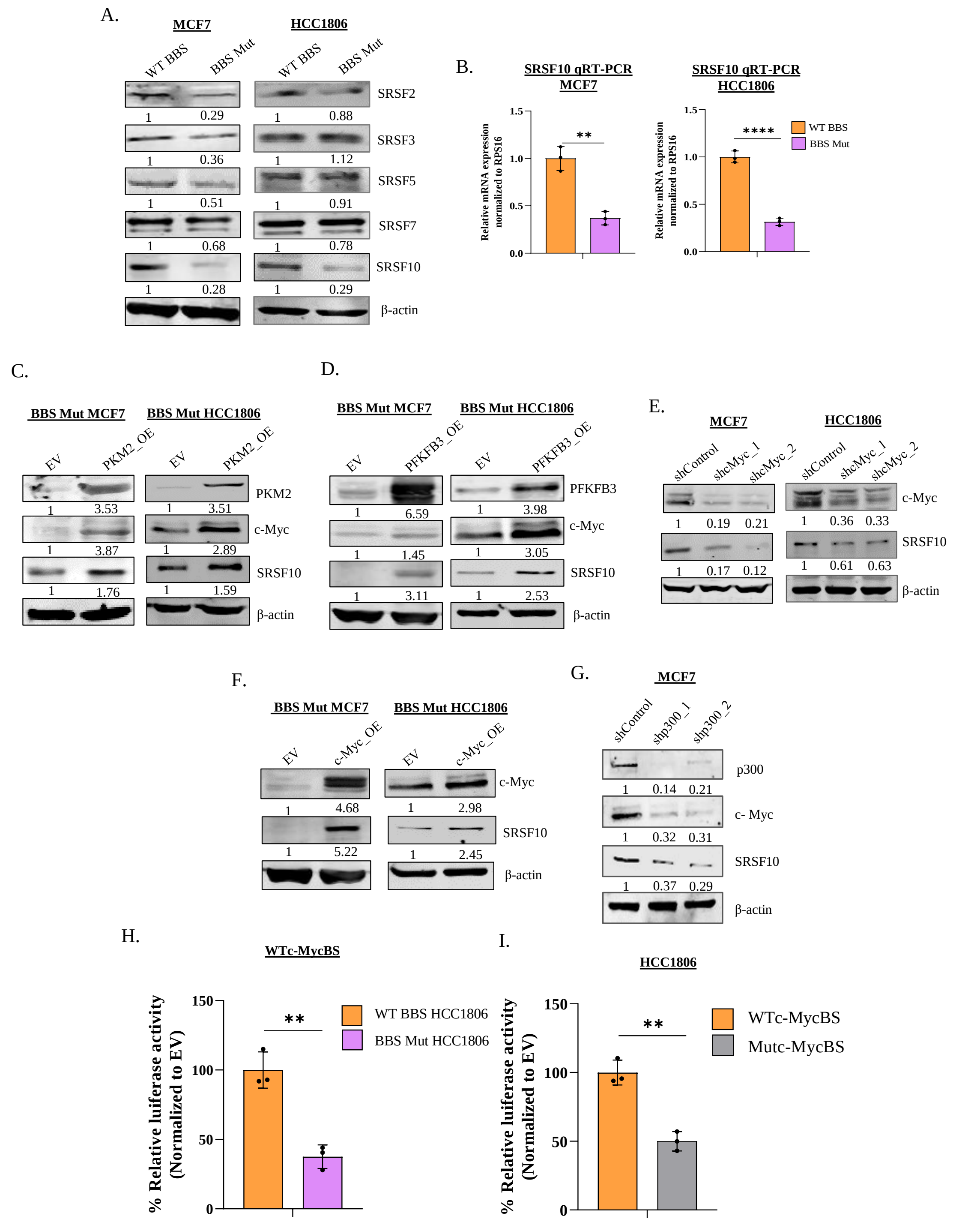


**Aerobic glycolysis favors SRSF10 expression.** *(A)* Immunoblot analysis of various SRSF family members performed in WT BBS and BBS Mut cell lines of MCF7 and HCC1806. *(B)* RPS16 normalized qRT-PCR analysis of *SRSF10* in WT BBS and BBS Mut cell lines. Immunoblot assay performed upon overexpressing *(C)* PKM2 and *(D)* PFKFB3 in BBS Mut cell lines. *(E)* Immunoblot analysis demonstrating the effect of c-Myc knockdown on SRSF10 expression in WT BBS cells. *(F)* Immunoblot assay performed upon overexpressing c-Myc in

BBS Mut cells. *(G)* Immunoblot analysis demonstrating the effect of p300 knockdown on c-Myc and SRSF10 expression in WT BBS MCF7 cells. *(H)* Luciferase assay of WTc-MycBS SRSF10 promoter construct performed in WT BBS and BBS Mut HCC1806 cells. *(I)* Luciferase assay of WTc-MycBS and Mutc-MycBS SRSF10 promoter constructs performed in WT BBS HCC1806 cells. Data are represented as mean± SD. *$P \leq 0.05$.

### Supporting information 4 (S4)

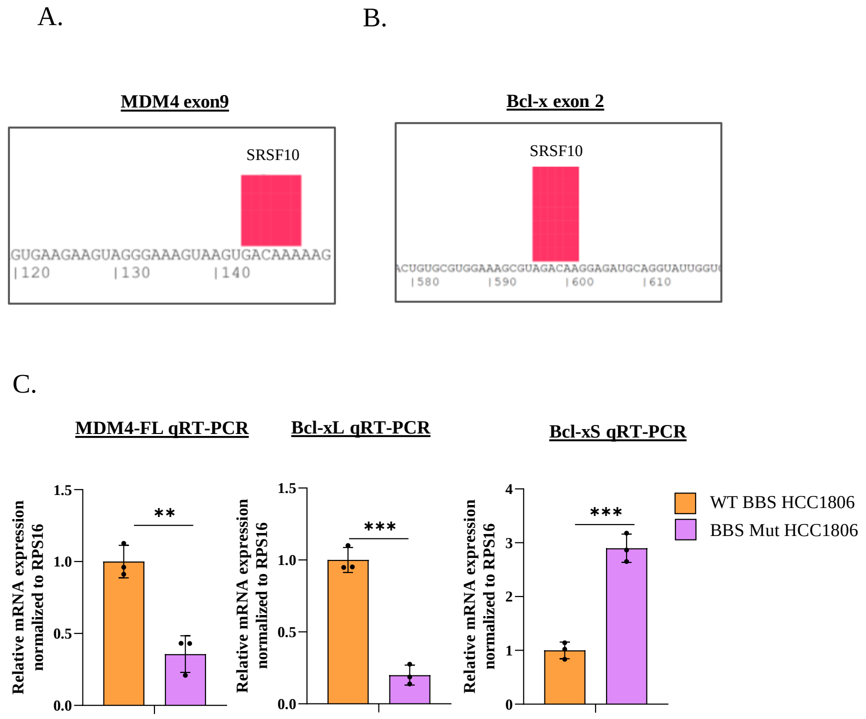


**SRSF10 binds to MDM4 and Bcl-x mRNA.** *(A)* Predicted SRSF10 binding site on exon 9 of MDM4. *(B)* Predicted SRSF10 binding site on exon 2 of Bcl-x. *(C)* RPS16 normalized qRT-PCR analysis of MDM4-FL, Bcl-xL and Bcl-xS in WT BBS and BBS Mut HCC1806 cells. Data are represented as mean± SD. *P ≤ 0.05.

### Supporting information 5 (S5)

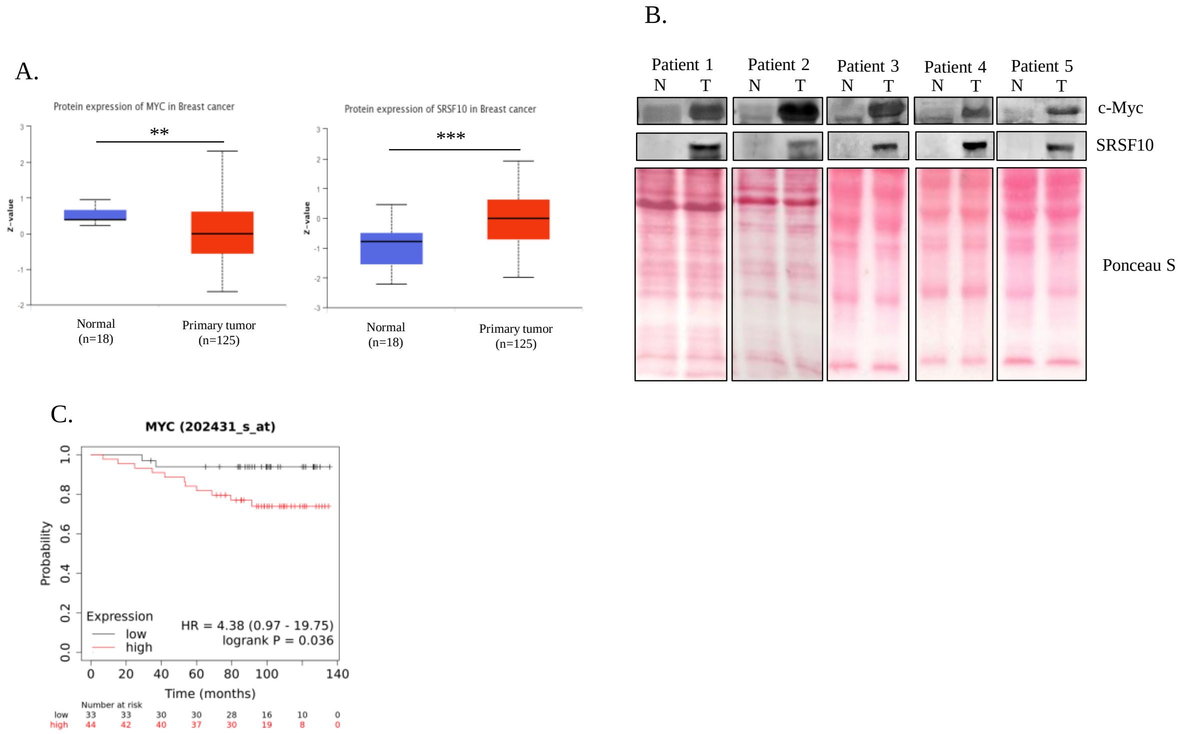


**High c-Myc expression is associated with poor breast cancer prognosis.** *(A)* Protein expression profile of c-Myc and SRSF10 in normal and breast cancer samples obtained from CPTAC platform. *(B)* Immunoblotting analysis showing c-Myc and SRSF10 expression in 5 representative breast cancer patient samples. *(C)* Kaplan–Meier curve showing significant association (p-value 0.036) of Recurrence Free survival with c-Myc expression in Breast Cancer TCGA dataset (GSE9195). $^*P \leq 0.05$.

### Supporting information 6 (S6)

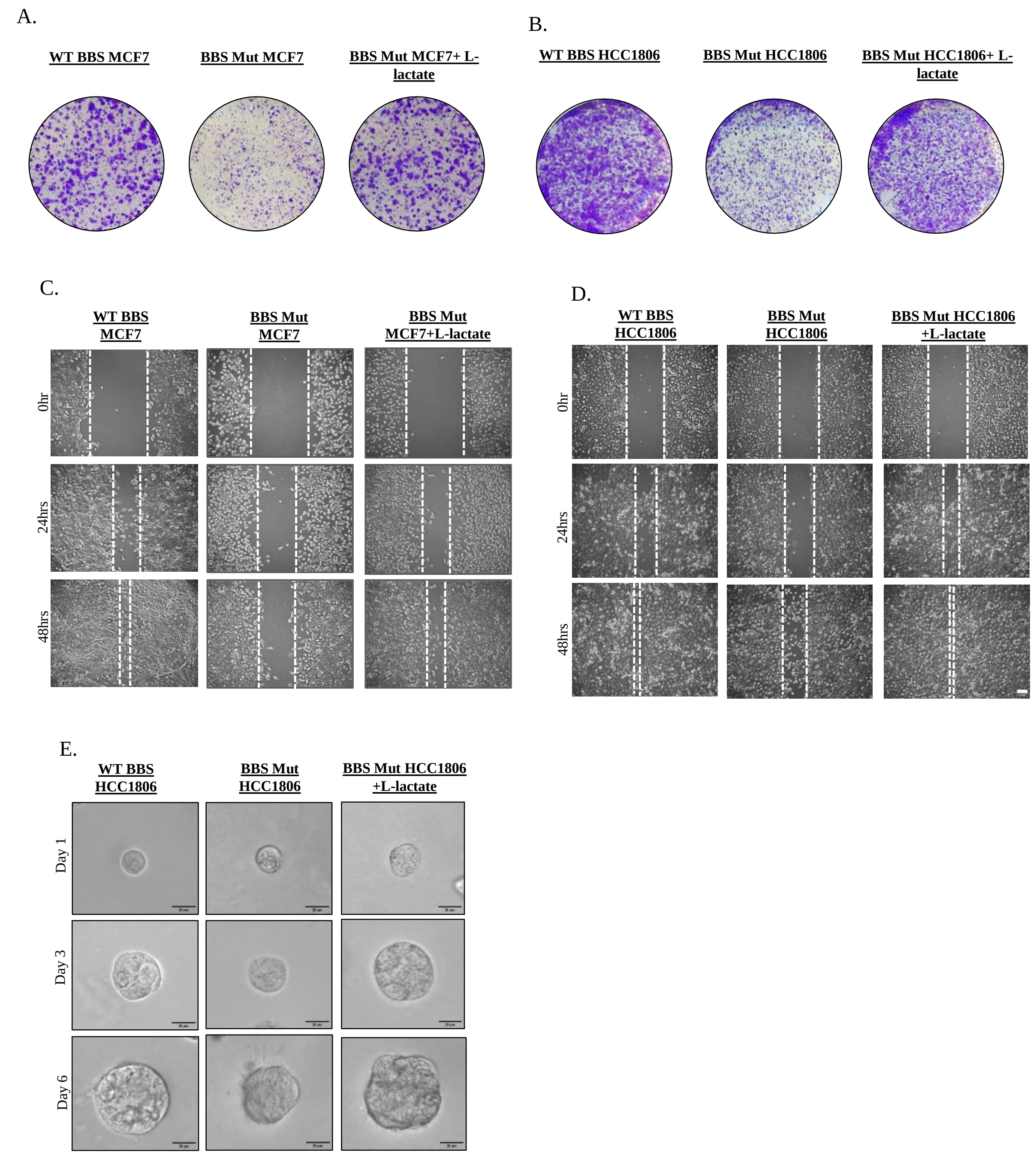


**c-Myc-SRSF10 axis promotes proliferation of breast cancer cells.** Colony formation assay performed in *(A)* WT BBS, BBS Mut and BBS Mut MCF7 cells subjected to 5mM L-lactate treatment for 10 days and *(B)* WT BBS, BBS Mut and BBS Mut HCC1806 cells subjected to 5mM L-lactate treatment for 10 days. Cell migration analysed via wound healing assay in *(C)* WT BBS, BBS Mut and BBS Mut MCF7 cells subjected to 5mM L-lactate treatment for 48hrs and *(D)* WT BBS, BBS Mut and BBS Mut HCC1806 cells subjected to 5mM L-lactate treatment for 48hrs. *(E)* Spheroid formation assay performed in WT BBS, BBS Mut, and BBS Mut MCF7 cells subjected to 5mM L-lactate treatment for 6 days.

**Table S1: List of antibodies**

| Antibody | Company | Catalogue no. | Lot no. |
|---|---|---|---|
| PKM2 | Cell Signalling Technology | 4053S | 5 |
| Anti-PFKFB3 | Abcam | ab181861 | GR3256208-2 |
| Alexa-Flour 680 anti-rabbit IgG | Invitrogen | A32734 | RJ243414 |
| Alexa-Flour 800 anti-mouse IgG | Invitrogen | A32730 | SC243837 |
| Normal Rabbit IgG | Cell Signalling Technology | 2729S | 8 |
| P300 | Cell Signalling Technology | 54062S | 1 |
| Anti SRSF10 Antibody | Sigma | HPA053831-100ul | 000002200 |
| β-Actin (D6A8) Rabbit mAb | Cell Signalling Technology | 8457S | 7 |
| c-Myc | Cell Signalling Technology | 9402S | 11 |
| DYKDDDDK Epitope Tag Ab (L5) (flag) | Novus | NBP1-06712SS | C-4 |
| Lactyl-Histone H3 (Lys18) Rabbit Monoclonal Antibody | PTM Biolabs | PTM-1406RM | RM021106 |
| SRSF2 | Sigma | HPA049905-100ul | 000015636 |
| SRSF3 | Abcam | ab73891 | GR321343-12 |
| SRSF5 | Abcam | ab67175 | GR32337921-1 |
| SRSF7 | Abcam | ab170679 | GR179666-4 |

**Table S2: List of primers used for cloning**

| Primer name | Sequence (5'-3') |
|---|---|
| PKM2 OE Fw | TTTTGCGGCCGCATGTCACCGGAAGCCCAA |
| PKM2 OE Rv | TTTTGAATTCTCACGGCACAGGAACAAC |
| PFKFB3 NotI_Fw | TTTTGCGGCCGCATGCCGTTGGAACTGACGCAGA |
| PFKFB3 EcoRI_Rv | TTTTGAATTCTCAGTGTTTCCTGGAGGAGTCAGC |
| c-Myc OE_Fw | ATATGGATCCATGCCCCTCAACGTTAGCTTCAC |
| c-Myc OE Rv | ATATGAATTCTTACGCACAAGAGTTCCGTAGCT |
| SRSF10 Luc Fw | TTTTGGTACCAGAAAGAACAGGCTTGGGGGAG |
| SRSF10 Luc Rv | TTTTAAGCTTCAACGGTTCTTTCCCGCGAGAA |
| SRSF10 SDM Fw | GAATGACACCGACGCTGAGATATGTGCGCCAGCCCTCCGCATG |
| SRSF10 SDM Rv | CATGCGGAGGGCTGGCGCACATATCTCAGCGTCGGTGTCATTC |
| Myc -997 Fw | TTTTGGTACCGGAACAGGCAGACACATCTCAG |
| Myc -621 Fw | TTTTGGTACCGACTCTTGATCAAAGCGCGG |
| Myc -245 Fw | TTTTGGTACCGGCGTGGGGGAAAAGAAAAAAG |
| Myc +3 Fw | TTTTGGTACCCGCTGTAGTAATTCCAGCGAG |
| Myc +174 Rv | TTTTAAGCTTTTTCGTGGATGCGGCAAGGGTTG |

**Table S3: List of primers used for qRT-PCR**

| Primer name | Primer sequence (5’-3’) |
|---|---|
| RPS16 Fw | AAACGCGGCAATGGTCTCATCAAG |
| RPS16 Rv | TGGAGATGGACTGACGGATAGCAT |
| c-Myc Fw | TCCACCTCCAGCTTGTA |
| c-Myc Rv | TCGAGGAGAGCAGAGAAT |
| SRSF10 Fw | GGAGGAGATCAAGAAGTCGGTC |
| SRSF10 Rv | TGTCGGAATGGCTTCTGCTACG |
| SRSF10 ChIPFw | AGAAGCCAACGAATGACACC |
| SRSF10 ChIPRv | CTACCTCCGCGCAAAGTC |
| MDM4_8/9_Fw | ACAGACAAATCAGGATGTGGGTACT |
| MDM4_9_Rv | CCAACACCTAACTGCTCTGATACTG |
| Bcl-xS 2-3Fw | TATCAGAGCTTTGAACAGGATAC |
| Bcl-xS 3Rv | AAGAGTGAGCCCAGCA |
| Bcl-xL 2Fw | TCCTTCGGCGGGGCAC |
| Bcl-xL 2Rv | ACCCAGCCGCCGTTCT |
| Myc ChIP -245 Fw | GGCGTGGGGGAAAAGAAAAAAG |
| Myc ChIP -70Rv | GTTCTTTTTCCCGCCAAGCC |
| Myc ChIP -70 Fw | GGCTTGGCGGGAAAAAGAAC |
| Myc ChIP +3 Rv | CTCGCTGGAATTACTACAGCG |

**Table S4: Sequence of shRNA**

| Name | Sequence (5’-3’) |
|---|---|
| sh_Control | CCGGCGTGATCTTCACCGACAAGATCTCGAGATCTTGTCGGTGAAGATCACGTTTTT |
| shc-Myc_1 | CCGGCCTGAGACAGATCAGCAACAACTCGAGTTGTTGCTGATCTGTCTCAGGTTTTTG |
| shc-Myc_2 | CCGGCAGGAACTATGACCTCGACTACTCGAGTAGTCGAGGTCATAGTTCCTGTTTTTG |
| shSRSF10_1 | CCGGCCAGTACAGTTCTGCTTACTACTCGAGTAGTAAGCAGAACTGTACTGGTTTTTG |
| shSRSF10_2 | CCGGGCTATGATGATTATGACAGATCTCGAGATCTGTCATAATCATCATAGCTTTTTG |
| shp300_1 | CCGGCCTCACTTTATGGAAGAGTTACTCGAGTAACTCTTCCATAAAGTGAGGTTTTTG |
| shp300_2 | CCGGGCCTTCACAATTCCGAGACATCTCGAGATGTCTCGGAATTGTGAAGGCTTTTTG |

**Table S5: List of transcription factors and epigenetic factors differentially expressed in BBS Mut MCF7 vs. WT BBS MCF7 HTA 2.0 array**

| **Genes** | **Fold Change** |
|---|---|
| SMAD6 | -4.86 |
| MYC | -3.19 |
| FOXM1 | -3.81 |
| EGR1 | -2.55 |
| ARNT | -2.37 |
| E2F8 | -2.24 |
| ELF5 | -2.21 |
| SOX4 | 2.05 |
| STAT1 | 2.08 |
| STAT3 | 2.31 |
| EGLN3 | 2.97 |
| SOX9 | 4.29 |
| HIF1A | 5.73 |
| EPAS1 | 20.6 |
| TET2 | -5.95 |
| METTL7A | -4.46 |
| SMARCC2 | -3.07 |
| SETDB1 | -2.95 |
| SMYD4 | -2.83 |
| PRMT9 | -2.79 |
| TCFL5 | -2.63 |
| HLTF | -2.57 |
| YEATS4 | -2.36 |
| TCF7L2 | -2.21 |
| TEAD4 | -2.07 |
| SETD5 | -2.06 |
| SMARCA5 | -2.03 |
| MBD2 | -2.01 |
| METTL14 | -2.01 |
| SMARCD2 | 2.27 |

**Table S6: Details of the patient samples**

| S. No | Histopathology |
|---|---|
| 1 | Infiltrating duct carcinoma Grade II pT2N0Mx |
| 2 | Infiltrating duct carcinoma Grade1 pT1N0Mx |
| 3 | Infiltrating duct carcinoma Grade II pT2N0Mx |
| 4 | Infiltrating duct carcinoma Grade II pT2N1Mx |
| 5 | Infiltrating duct carcinoma pT1N0Mx |
| 6 | Residual tumor seen in a k/c/o carcinoma post NACT stage ypTmN1Mx (Considering limitations of metastasis) |
| 7 | Infiltrating duct carcinoma grade I Pathological stage pT(m)1N0Mx |
| 8 | Infiltrating duct carcinoma stage pT2N2Mx |
| 9 | Infiltrating duct carcinoma pT2NMx |
| 10 | Infiltrating duct carcinoma Grade II pT2N1Mx |
| 11 | Infiltrating duct carcinoma pT2N2Mx |
| 12 | Infiltrating duct carcinoma Grade I pT2N0Mx |
| 13 | Infiltrating duct carcinoma grade II Pathological stage pT4NxMx |
| 14 | Infiltrating duct carcinoma pT2N1Mx |
| 15 | Infiltrating duct carcinoma pT2N3Mx |
